# First report of overwintering tadpoles in the endemic Italian agile frog *Rana latastei*

**DOI:** 10.1101/2025.04.02.646811

**Authors:** Andrea Melotto, Antonio Serafino Melotto, Gentile Francesco Ficetola, Raoul Manenti, Mattia Falaschi

## Abstract

Climate change is affecting organism seasonality patterns, and this can drive phenological shifts for key traits, including breeding activity. Here we report a first record of overwintering tadpoles of the Italian agile frog (*Rana latastei*), a threatened endemic species of Northern Italy. This species usually breeds between late January and April, with tadpoles reaching metamorphosis by early summer. In February 2025, together with the first freshly-laid egg-clutches of the season, several large-sized tadpoles were found in one reproductive site of *R. latastei* in Como province. A few days after hatching, six individuals from newly hatched tadpole cohorts and six individuals from large-sized cohorts were captured, photographed and measured. Concurrently, a newly metamorphosed froglet was found. Results from morphological inspections revealed that individuals showed typical traits of *R. latastei*. Moreover, while newly hatched tadpoles were at development Gosner’s stages 25-26, large cohort individuals were visibly bigger and at late development stage, which was not compatible with the classical late winter breeding timing of this species. Our study shows for the first time a case of overwintering in *R. latastei*, suggesting shifts in this frog development trajectory (prolonged larval time) or breeding habits (autumnal reproduction) may be occurring. While mechanisms behind this unusual observation have to be ascertained, such a phenological shift might have been favoured by variation in climatic regime.

---

Global warming is causing unprecedentedly quick phenological shifts in animal populations (Horton et al., 2020). These shifts can involve breeding activities, including both anticipations (Parmesan, 2007; Ficetola and Maiorano, 2016) and delays (Dalpasso et al., 2023) of the start of the breeding season after the winter. Additionally, given the general increase of winter temperatures, the end of the activity season in autumn is expected to be postponed (Lang et al., 2025). This may affect voltinism, potentially opening new opportunities for increasing the number of broods per year in some populations, but can also produce survival risks for early stages, when suitable conditions for development are not matched (Benard, 2015; Bison et al., 2021). For instance, in temperate regions several amphibian species have been shown to extend or shift their breeding season by days or weeks in the last decades (Todd et al., 2011), while reports of amphibian or reptile species reproducing out of their typical breeding season exist (Graña and Martínez-Freiría, 2020; Rodriguez-Muñozet al., 2020).

In Europe, most anurans breed in late winter-spring, with juveniles metamorphosing in late spring-summer. This allows them exploiting the spring rain that fills temporary wetlands and growing through the warm season. However, breeding patterns can be different in some species of Southern Europe. For instance, in Sicily, the green toad (*Bufo boulengeri*) can also breed in autumn and winter, with tadpoles from autumnal mating overwintering and metamorphosing in early spring (Sicilia et al., 2006). In this case, autumnal breeding has been interpreted as an adaptation towards dry environments, as wetlands filled by autumn rain have a higher probability to retain water long enough to enable toadlets attaining metamorphosis (Sicilia et al., 2006). Less evidence of autumnal breeding and tadpole overwintering is available for amphibians living in Northern Italy. For instance, the Italian Agile frog *Rana latastei*, is endemic of the lowlands of Northern Italy and adjacent areas and breeds from late January to mid-April (Ambrogio and Mezzadri, 2018; Ficetola et al., in press). In principle, changes in temperature and/or precipitation patterns related to climate change might affect breeding phenology of this species, but information on autumn breeding and/or overwintering tadpoles is so far lacking.

Here, we report for the first time a case of overwintering tadpoles for the Italian agile frog. The reproductive site is a small artificial permanent pond (roughly 2 × 1 m; 0.3 m depth) located in a small wooded area in the foothills of the Como district (Inverigo, Lombardy, Italy). The site is located approximately 350 m a.s.l. and is home to numerous water bodies that support a diverse amphibian community, including three urodeles and six anuran species. This area has been the subject of long-term amphibian monitoring that has spanned the past two decades (Ficetola et al., 2009; Falaschi et al., 2021), which revealed a fairly stable meta-population for *R. latastei* among interconnected reproductive sites (Manenti et al., 2020). The permanent pond where overwintering tadpoles were found has been consistently surveyed multiple times during spring since 2010 and is a stable breeding site of *R. latastei*, where no other amphibian, except for salamanders, has been reported reproducing. On 27^th^ January 2025, ~5 days after mid-winter rainfalls that usually give start to the amphibian breeding season in the study area, the site was monitored along with other sites in the surroundings. This survey revealed the presence of a single freshly laid egg-clutch of *R. latastei* (the first clutch of the breeding season found in the area). The site was then monitored on January 30 (1 new clutch), February 6 (no new clutches), February 11 (1 new clutch), and on February 26 (4 new clutches). The first egg hatch was observed on February 11. On February 6, the presence of three large-size tadpoles showing *R. latastei* traits was observed. On February 26, we collected 6 large-sized tadpoles and 6 small-sized tadpoles (Gosner’s stage >25) for measuring (Fig. 1). Contextually, a newly metamorphosed froglet was found (Fig. 2). Individuals were captured by gently netting the pool then briefly kept in a small plastic tank and photographed on graph paper using a 100 mm macro lens ensuring limited distortion. After the photoshoot, the individuals were immediately released (permits in acknowledgements). Froglet and tadpoles were measured from scaled pictures using ImageJ software (Schneider et al., 2012) following standard procedures (Relyea, 2001; Melotto et al., 2020;

**Figure 1.**
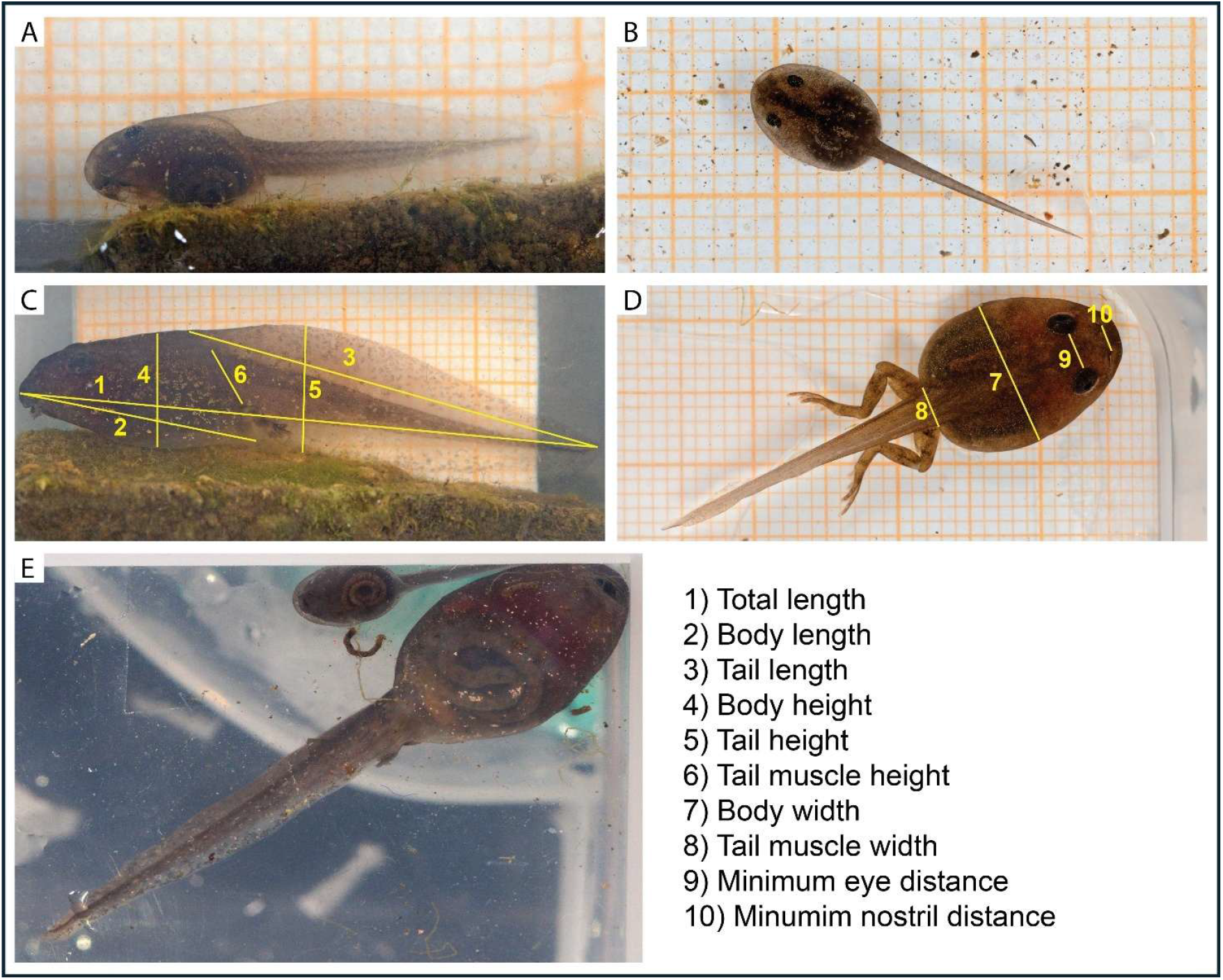
Tadpoles of *Rana latastei*. A-B) Lateral and dorsal view of newly hatched individuals; C-D) lateral view of overwintering individuals; E) ventral view comparison between a newly hatched and an overwintering individual. For C e D, examples of measures performed are provided.

**Figure 2.**
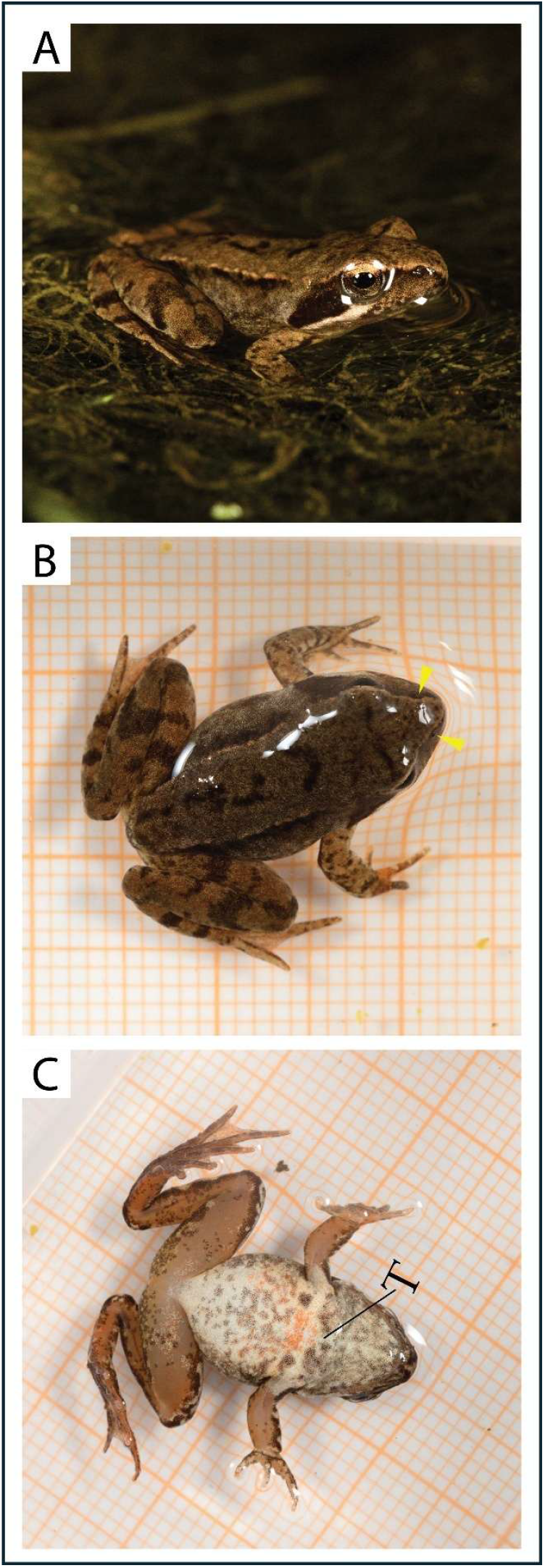
Newly metamorphosed *Rana latastei* individual. A) lateral view, white stripe stopping under the eye visible; B) dorsal view, nostrils are highlighted by yellow arrow-heads; C) ventral view, T shape between throat and forelimb junction highlighted.

Melotto et al., 2021). Tadpole development stages were determined basing on Gosner’s tables (Gosner, 1960). Even if never observed breeding in the study pond, another *Rana* species with similar ecology, the agile frog (*Rana dalmatina*), is present in the nearby woods, and tadpoles of this frog sometimes show phenotypes difficult to tease apart from *R. latastei* (Barbieri et al., 2000). Hereafter, along with morphometric and stage comparison between tadpole cohorts, key phenotypic traits differing between the two frog species are highlighted (Lanza et al., 2009; Ambrogio and Mezzadri, 2014).

We obtained tadpole’s measurements from both dorsal and side pictures (Table 1A, Fig. 1A-D) and included total length (Totl), body length (BL), body height (BL), body width (BW), tail length (TL), tail height (TH), tail muscle height (TMH), and tail muscle width (TMW). Moreover, we used dorsal pictures to calculate minimum eye distance (ED) and minimum nostril distance (ND). ND was available for large size tadpole only, as nostril position was not clearly identifiable in newly hatched tadpoles. All tadpole measurements, except for ND, were included in a principal component analysis to ascertain the existence of distinct size classes among tadpoles from different cohorts and if they matched the eye-based and Gosner’s stage identification. Moreover, we calculated eye-distance nostril-distance ratio (ENDr) from ED and ND measurements for the perspective overwintering tadpoles (large size cohort individuals). This ratio is one of the key traits differing in tadpoles of *R. dalmatina* and *R. latastei*, being around 2 for the first one and rarely >1.5 for the latter (Lanza et al., 2009). Finally, we used ventral pictures of the froglet to take some morphological measurement (Table 1B): total length (TL), body width (BW), jaw width (JW), and few left hindlimb lengths, including proximal hindlimb (LPHL), distal hindlimb (LDHL), tarsus (LTL), and foot (LFL). Additionally, froglet ED and ND were obtained from dorsal picture (Table 1B).

**Table 1.** Morphological assessment. A) measures and approximated Gosner’s development stage of tadpoles of different cohorts are reported together with identity of the individual, size class of the cohort; B) measures of the newly metamorphosed froglet found. Measurements abbreviations hereafter: body height (BH), tail height (TH), tail muscle height (TMH), body length (BL), tail length (TL), total length (TotL), body width (BW), tail muscle width (TMW), eye distance (ED), nostril distance (N D), eye-nostril distance ratio, jaw width (JW), left distal hindlimb length (LDHL), left proximal hindlimb length (LPHL), left tarsus length (LTL), left foot length (LFL).

| A) | ID | cohort | Gosner | BH | TH | TMH | BL | TL | TotL | BW | TMW | ED | ND |
| --- | --- | --- | --- | --- | --- | --- | --- | --- | --- | --- | --- | --- | --- |
|  | GIR1 | large | 37 - 39 | 0.884 | 1.042 | 0.510 | 1.858 | 3.149 | 4.525 | 1.089 | 0.349 | 0.341 | 0.244 |
|  | GIR2 | large | 40 | 0.912 | 0.810 | 0.405 | 1.563 | 2.320 | 3.489 | 1.049 | 0.301 | 0.267 | 0.232 |
|  | GIR3 | large | 40 | 0.811 | 0.858 | 0.403 | 1.539 | 2.450 | 3.626 | 1.027 | 0.289 | 0.263 | 0.220 |
|  | GIR4 | large | 40 | 0.847 | 0.826 | 0.375 | 1.550 | 2.347 | 3.467 | 0.917 | 0.363 | 0.274 | 0.192 |
|  | GIR5 | large | 40 | 0.857 | 0.983 | 0.423 | 1.563 | 2.461 | 3.698 | 1.076 | 0.312 | 0.323 | 0.247 |
|  | GIR6 | large | 37 - 39 | 0.992 | 1.069 | 0.606 | 1.850 | 3.483 | 4.841 | 1.136 | 0.379 | 0.332 | 0.259 |
|  | GIR7 | small | 25 - 26 | 0.374 | 0.423 | 0.178 | 0.633 | 1.285 | 1.772 | 0.395 | 0.128 | 0.119 | - |
|  | GIR8 | small | 25 - 26 | 0.370 | 0.403 | 0.172 | 0.624 | 1.194 | 1.685 | 0.404 | 0.091 | 0.138 | - |
|  | GIR9 | small | 25 - 26 | 0.347 | 0.387 | 0.168 | 0.662 | 1.206 | 1.740 | 0.455 | 0.095 | 0.157 | - |
|  | GIR10 | small | 25 - 26 | 0.362 | 0.404 | 0.167 | 0.683 | 1.258 | 1.696 | 0.431 | 0.091 | 0.146 | - |
|  | GIR11 | small | 25 - 26 | 0.297 | 0.344 | 0.166 | 0.582 | 1.201 | 1.585 | 0.367 | 0.091 | 0.128 | - |
|  | GIR12 | small | 25 - 26 | 0.361 | 0.404 | 0.171 | 0.700 | 1.299 | 1.781 | 0.404 | 0.082 | 0.126 | - |
| B) | ID | TL | BW | JW | LDHL | LPHL | LTL | LFL | ED | ND | ENDr |  |  |
|  | froglet | 1.867 | 0.888 | 0.819 | 0.968 | 0.952 | 0.480 | 0.899 | 0.437 | 0.195 | 2.241 |  |  |

Overall, all tadpoles showed characters matching the typical features of *R. latastei* [absent, poor, or incomplete ventral colouration and visible guts (Fig. 1E); EDNr of large-sized tadpoles ranged from 1.15 to 1.43 (average: 1.29) (Lanza et al., 2009; Ambrogio and Mezzadri, 2014)]. All the small-sized individuals were between Gosner’s stage 25 and 26, showing functional oral canal but no trace of hindlimb formation. Instead, large size-tadpoles were at the Gosner’s stage 40 (four individuals; hindlimb with clearly differentiated toes and tubercles, cloacal tail piece visible), or between 37 and 39 (evident toes with no tubercles). The principal component analysis revealed that the first axis explained 99% of variation, identifying three distinct size classes corresponding to the three Gosner’s stage identified, clearly distinguishing as different classes large size individuals and newly hatched ones (Fig. 3), also evident from picture comparison (Fig. 1E).

**Figure 3.**
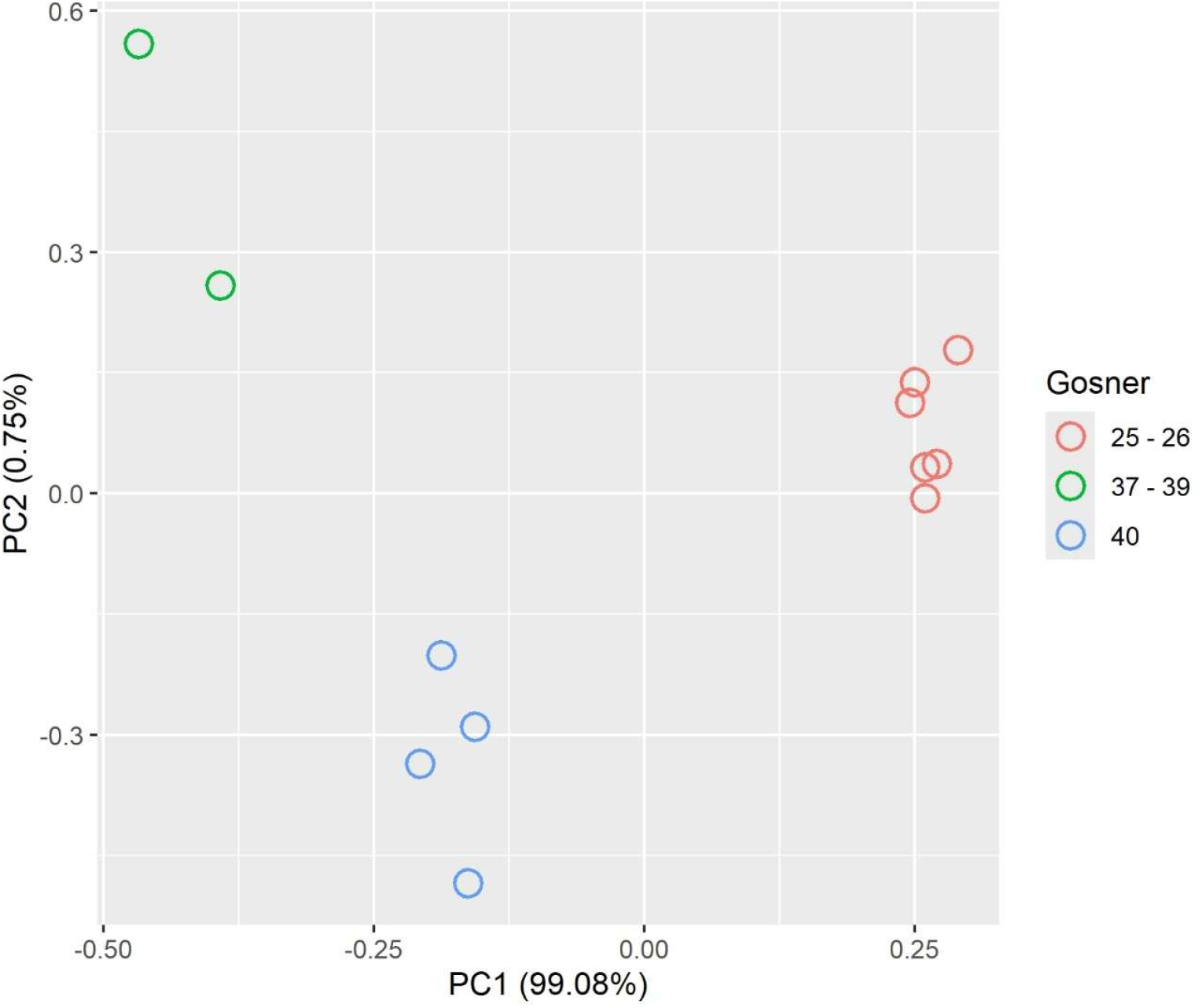
Principal component analysis performed using Tadpole measures in Table 1A, excluding “ND” and “ENDr”, which were not measured for Gosner’s 25 – 26 tadpoles. Colours represent Gosner’s developmental stages, as indicated in Table 1A.

The newly metamorphosed individual measured 1.87 cm (Table 1B) and presented the typical traits of *R. latastei* (Lanza et al., 2009): the light stripe running above upper lip suddenly stops under the eye (while it’s generally prolonged in *R. dalmatina*), and nostril distance was shorter than eye distance (Fig. 2A-B, Table 1B); moreover, chest and throat looked diffusely spotted and a “T” shaped stripe is noticeable between the throat and the forelimb junction (Fig. 2C), while ventral colouration is generally homogeneously pale and unspotted in *R. dalmatina*.

Taken together, our observations confirm that all the individuals belong to *R. latastei*. At the same time, the size and late stage of the large tadpoles and the presence of a froglet, along with the comparison with newly hatched tadpoles, suggest that large-size individuals originated from reproductive events occurred in the previous year and underwent overwintering at larval stage. In absence of a direct observation of autumnal reproductive events, only speculations can be made regarding the breeding period. Overwintering tadpoles are not rare in anurans (McDiarmid and Altig, 1999) and occasionally occur in other local species (for instance green frogs, *Pelophylax* synkl. *esculentus*) and some *Rana* species in other areas (Walsh et al., 2008; Lanza et al., 2009). However, this is generally observed in anurans facing shorter growing seasons, such late breeding species, or species and populations from high latitude or elevation (McDiarmid and Altig, 1999), as overwintering can extend growth period and allow attaining larger size at metamorphosis (Walsh et al., 2008; Iwai, 2024). This is not the case of *R. latastei*, which is an early breeder, whose tadpoles typically reach metamorphosis in June-July (Lanza et al., 2009).

We suggest that these tadpoles originate from one or more breeding events occurring in late summer-autumn 2024. Tadpoles exposed to low temperatures typically incur in metabolic depression, which slows down growth and development (McDiarmid and Altig, 1999; Enriquez-Urzelai et al., 2022). However, autumn and winter temperatures of 2024-2025 have been among the mildest of the last decades and 2024 has been the warmest year since temperature has been consistently recorded (https://climate.copernicus.eu/). This could have allowed larval development during the winter months, similarly to what happens in anurans breeding in warmer regions (Sicilia et al., 2006). Additionally, previous records of *R. latastei* males calling in autumn have been already reported for nearby areas (Grossenbacher et al., 2000). While the presence of calling males does not guarantee that breeding activities are occurring, milder autumn and winter temperatures may encourage egg deposition and allow tadpole development during this time.

An alternative hypothesis might be that large cohort tadpoles originated from a typical late-winter deposition event in spring 2024. Breeding site monitored is partly shaded by canopy cover, and 20 egg-clutches were laid by February 2024 (last deposition event observed). We cannot exclude that tadpole density and cold water temperatures might have induced delayed development in some individuals at the breeding site. However, tadpole density was comparable to other nearby sites monitored where *R. latastei* metamorphosed as usually in early summer. Moreover, June-September temperatures of 2024 (mean air temperature ± SD: 29.7 ± 5.1 °C) were visibly higher than those experienced over the typical growing season (March-June temperatures: 21.4 ± 6.1), making it extremely unlikely that overwintering individuals were tadpoles originating from late-winter 2024 deposition and then undergoing delayed development because of low temperature.

Whatever the period of deposition, the present report highlights the existence of an unusual shift in breeding phenology of *R. latastei*, which calls for further investigations assessing the frequency of these events and unravelling drivers and potential implications of such a shift. Indeed, variation in reproductive timing can result in crucial consequences for individual life-history and survival (Bison et al., 2021; Enriquez-Urzelai et al., 2022); moreover, in the case of massive, explosive breeders, like the wood frogs, this can also produce cascading effects on amphibian populations and freshwater community dynamics, with complex outcomes difficult to foresee (Todd et al., 2011). This first record of overwintering tadpoles in the Italian agile frog may represent an anomalous or isolated event, but correlation between increasing temperatures, changes of precipitation patterns and shifts in amphibian breeding period have been observed in multiple species as a strategy to withstand global warming (Todd et al., 2011; Ficetola and Maiorano, 2016). Under the ongoing climate change scenarios, amphibian phenology is expected to be considerably impacted worldwide, and variations in their reproductive activity should deserve particular attention. As for the case here reported, autumnal monitoring of the study breeding sites and those nearby will be planned during the incoming years to ascertain the occurrence of *R. latastei* depositions outside the typical reproductive season.

